# Seasonal and social factors associated with spacing in a wild territorial electric fish

**DOI:** 10.1101/2020.01.29.924761

**Authors:** Lucía Zubizarreta, Laura Quintana, Daniel Hernández, Franco Teixeira de Mello, Mariana Meerhoff, Renato Massaaki Honji, Renata Guimarães Moreira, Ana C. Silva

## Abstract

The expression of territorial behavior in wild species is especially suited to explore how animals integrate individual traits with dynamic environmental and social contexts. In this study, we focused on the seasonal variation of the determinants of territory size in the weakly electric fish *Gymnotus omarorum*. This species is a seasonal breeder that displays year-long territorial aggression, in which female and male dyads exhibit indistinguishable non-breeding territorial agonistic behavior and the only significant predictor of contest outcome is body size. We carried out field surveys across seasons that included the identification of individual location, measurements of water physico-chemical variables, characterization of individual morphometric and physiological traits, and their correlation to spatial distribution. Although *Gymnotus omarorum* tolerates a wide range of dissolved oxygen concentration, territory size correlated with dissolved oxygen in both seasons. In the non-breeding season, we show that territory size is sexually monomorphic and explained only by body size. In the breeding season, while body size no longer correlated with territory size, evidence of sexual differences in territory size determinants emerged. First, the overall spatial arrangement adopted a sexual bias. Second, territory size depended on gonadal hormones in both sexes, which was expected for males, but not previously reported in females. Third, females’ territory size correlated with gonadal size and females showed relatively larger territories than males, probably to meet sexually dimorphic energetic requirements. This study provides evidence of seasonal changes in factors correlated with territory size and contributes to the understanding of the mechanisms underlying behavioral plasticity.

## Introduction

The mechanisms underlying behavioral plasticity, by which animals adapt to dynamic environmental and social contexts, are far from being fully understood (1). The study of the modulation of territorial behavior in wild model species is especially suited for this aim, as animals assess the environmental and social clues that determine territory quality and this information is contrasted with individual requirements and fighting abilities, to decide whether to compete over an area or not. Therefore, the distribution of territorial animals in space provide a hint on the integration of individual traits with environmental and social factors. Variation in the ability or motivation to obtain and defend a territory can generate differences in territory size, as traits such as body mass, sex, and reproductive state are known to influence resource holding potential and resource value (2–6). Within a population, body size is associated with territory size across species, as it directly correlates with metabolic requirements. Body size is the universal indicator of physical strength and thus it strongly impacts on contest outcome and territory size (7–10). In species that display territoriality in both sexes, asymmetry in fighting abilities or motivational factors may lead to sex differences in territory size; for example, in red squirrels (*Sciurus vulgaris*), in which males often hold larger territories than females (11) or in the stripped plateau lizard (*Sceloporus virgatus*), in which females are more territorial than males (12).

Many species show territorial behavior only during the breeding season (13). On the other hand, some species across different phylogenetic groups, in spite of being seasonal breeders, show robust territorial aggression all year round, as has been reported in birds (14–17), mammals (18–20), reptiles (21), and fish (22,23). These species offer a valuable opportunity to study the seasonality of environmental features and individual traits, and their relation to territory size in the natural habitat. During the breeding season, male territorial aggression is largely dependent on sexual gonadal steroids across vertebrates (24–26) and, in particular, androgen levels have been related to territory size in the wild (27–30). In contrast, in breeding females, there are few studies on the association between estrogen (E_2_) circulating levels and territorial aggression in free-living conditions (31–33), and, to our knowledge, there are no studies reporting the association between circulating E_2_ and territory size.

The weakly electric South American fish, *Gymnotus omarorum* (34), is a seasonal breeder that displays male and female territorial aggression all year-long and thus is an interesting model system to study the seasonal control of territoriality and its sex differences. Previous laboratory results showed that this species presents a remarkably robust non-breeding territorial aggression (initially described in (22)), with well-characterized agonistic behavioral displays including modulations of the electric organ discharge (EOD) to signal submission (35–37) and a dominant phenotype that persists for at least 36 hours (38). Under experimental laboratory conditions, male-male and female-female dyads that display non-breeding territorial behavior have shown no differences in either contest outcome, temporal dynamics of the agonistic encounter, levels of aggression, nor submissive signaling (39). Moreover, the only significant predictor of contest outcome is body size (22), and none of the features of agonistic encounters depends on circulating gonadal hormones (40).

In this study, we aimed to evaluate the seasonal variation of the ecological, morphometric, and physiological correlates of territory size in the wild in *Gymnotus omarorum*. In the non-breeding season, when gonads are regressed and thus circulating gonadal hormones are low, we stand on previous behavioral results to predict that territory size would be sexually monomorphic and explained mostly by body size, regardless of variations in local environmental characteristics. In the breeding season, when motivational aspects of territoriality may be confounded with the reproductive drive, we expected the emergence of sexual dimorphism in territory size determinants.

## Materials and Methods

### Study location and sampling seasons

Fieldwork was carried out in the Laguna de los Cisnes, Uruguay (205 ha, 34° 48’ S, 55° 18’ W) which composes a three-part interconnected shallow system (maximum depth 5 m) and has no inputs of salt or brackish water (Fig. 1A, (41)). The study species, *Gymnotus omarorum*, is a weakly electric fish, the only one present in the area of study from the several electric species present in Uruguay. The littoral area of the lake is blanketed by a strip (5-40 m with) of dense mats of free-floating aquatic macrophytes that cover the sampling area (Fig. 1B), with *G. omarorum* typically living among the roots of these plants (34).

**Figure 1:**
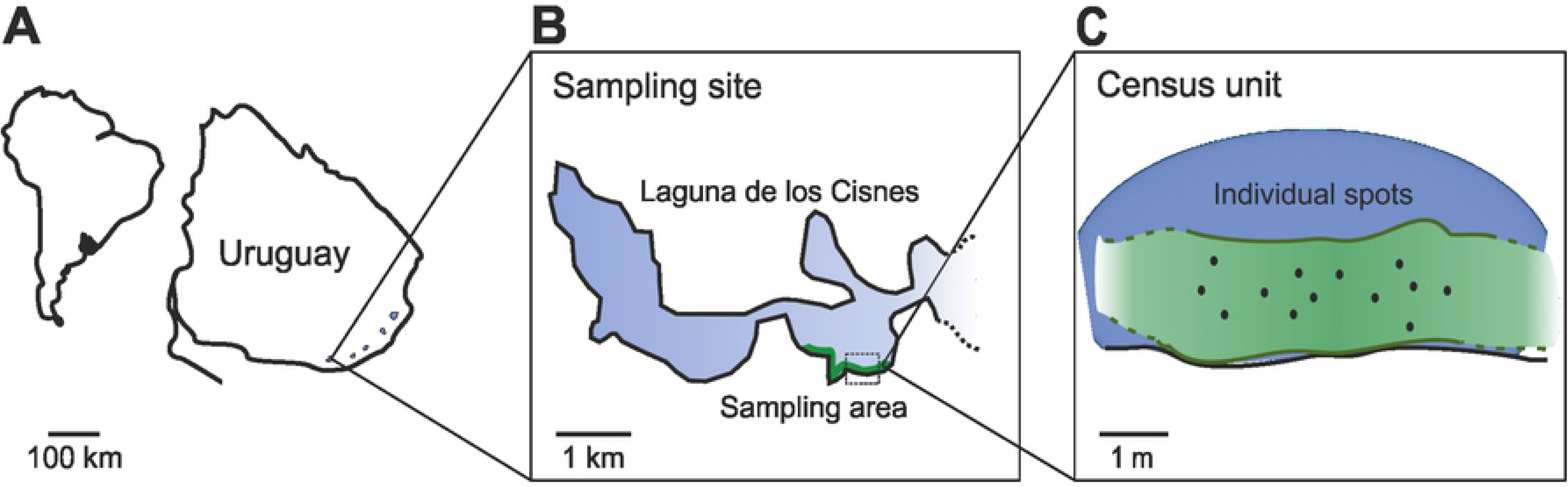
Study site and sampling method. A. The study site is located in Maldonado, Uruguay, in Laguna de los Cisnes. B. The shores of the lake have water hyacinths creating extensive floating mats that constitute the sampling area. C. Census unit illustrating individual spots. Fish location in individual spots was achieved by carrying out an electric census. Once a fish was located, water dissolved O_2_ concentration, temperature, and fish EOD rate were measured in each spot.

Following Quintana et al. (2004), who identified the breeding season for weakly electric fish species in the neotropical subtropical region from December to February (austral summer), we collected field data during December, corresponding to the early-breeding season before the appearance of offspring (42), and from June to August (austral winter), corresponding to the non-breeding season.

### Sampling method

Sampling area was homogeneous in depth, distance to shore, and vegetation composition. The sampling area was divided into adjacent transects referred to as census units (Fig. 1B and 1C, defined below), which were each studied in different days, without repeating sites. Sampling was performed during the day, which is the resting phase of animals in this species (43), in two stages: the first one (field survey 1) during the morning from 8 to 12 AM, and the second one (field survey 2) from 1 to 6 PM.

All research procedures complied with ASAP/ABS Guidelines for the Use of Animals in Research and were approved by the Institutional Ethical Committee (Comisión de Ética en el Uso de Animales, Instituto Clemente Estable, MEC, 02-2012).

<u>Field survey 1</u>: Electric census and environmental variables.

In order to achieve a first picture of the spatial arrangement of individuals, we carried out an electric census during the resting phase of the animals and measured the distance between each fish and its nearest neighbor (distance to the nearest neighbor, DNN) in the two seasons. To calculate DNN, we considered each focal fish distance to the nearest neighbor, even if the neighbor was not retrieved. The electric census implies the location of individual fish by means of an electronic audio amplifier connected to a pair of electrodes, as described elsewhere (44), and enables to locate individual fish through its EODs when the sound is maximal (detection range is 60 cm). The EOD rate monitoring also allows researchers to confirm that fish remain undisturbed during the measurement of parameters. Gymnotids show a typical behavior called the novelty response (reviewed in (45)), which consists of a transient increase in the EOD rate triggered by changes in nearby impedance, i.e., changes in its surrounding environment (46). To perform the electric census, two experienced researchers waded up to 1.2 m depth in the water, to slowly and carefully access places 1 m away from individual fish. Once a fish was located in an individual spot (Fig. 1C), the plant above it was tagged, and conductivity, dissolved oxygen concentration (O_2_, mg/l) and water temperature (T, °C) were measured 30 cm below the surface (using TDSu Testr 3, Cole Parmer for conductivity, and OxyWard, Handy Polaris for O_2_ and T). After taking physicochemical measurements, the EOD of each fish was recorded in situ for 10 seconds by means of two electrodes lowered in the vicinity of the animal, connected (by 10-30 m cables) to an amplifier located on the shore of the lake (World Precision Instruments Inc., Sarasota, FL. DAM-50, AC-coupled). After amplified, signals were recorded on a portable computer, captured by the audio card and stored for further analysis. In order to normalize the potential effect of water temperature on EOD rate, values were corrected to a constant 20°C temperature by using the Q10 value of 1.5 as calculated for electric fish (47,48). Q10 is a unitless quantity calculated as the factor by which the EOD rate increases when the temperature (T) is raised by 10 °C and is calculated as: Q10 = EOD rate × T / EOD rate × (T + 10).

During each sampling day, the measurement of physico-chemical water parameters and EOD rate was repeated for all the fish located within a census unit. A census unit was defined as the area with all fish detected until 12 AM, or the area where we detected a group of fish surrounded by at least 6 meters of water uninhabited by the species (Fig. 1C).

<u>Field survey 2</u>: Capture and quantification of individual traits.

In addition to environment variables, individual traits can influence spatial distribution and be different depending on the season. Therefore, based on field survey 1, we characterized morphometric and physiological traits of retrieved fish in both seasons, and then analyzed its correlation to spatial distribution. Individual spots were revisited in order, and each fish located under the tagged plants was collected using a net.

In the breeding season, immediately after netted, fish were anesthetized by immersion in a fast-acting eugenol solution (1.2 mg/l, first dissolved in alcohol 70%) for blood sampling from the caudal vein with a heparinized syringe in less than 3 min, which is the time range usually used to avoid a stress response due to manipulation (49–51). Captured fish were then weighed, measured and their gonads were visually inspected for sex determination. Blood was placed in tubes in ice to be centrifuged and stored at −80 °C in the laboratory (six hours later), gonads dissected in the field were stored in dry ice, and then weighed in the laboratory for gonadosomatic index (GSI) calculation. The index was later calculated for all adult animals as [Gonad Weight / Total Tissue Weight] × 100 (52).

### Hormone assays

Blood samples were taken in the field as described above, and once in the laboratory plasma was separated by centrifuging the samples at 3000 rpm for 10 min and stored at −80 °C until assayed. 17-β Estradiol (E_2_) levels were quantified in breeding females, and 11-Ketotestosterone (11-KT) in breeding males by enzyme-linked immunosorbent assay (ELISA) using commercial kits (IBL International, Hamburg, Germany for E_2_ and Cayman Chemical Company, MI, USA for 11-KT). The analyses were carried out according to the manufacturer’s instructions and a standard curve was run for each ELISA plate. In all cases, samples were assayed in duplicate and analyses were carried out on samples whose coefficients of variation were below 20% (53). Intra-assay variation was 3.95 % for E_2_ (detection limit: 25 pg/ml) and 6.2 % for 11-KT (detection limit: 1.56 pg/ml). Pilot assays using three different dilutions of 8 samples (4 samples per sex) were run to establish the appropriate working dilutions, which were 1:2 for breeding female E_2_ and 1:30 for breeding male 11-KT. The assays were validated with standards provided in the kit, indicating that each assay effectively detects *G. omarorum* E_2_ and 11-KT.

### Data analysis and statistics

All data were subjected first to D’Agostino & Pearson omnibus normality test. If data fitted to a gaussian distribution, were analyzed with parametric tests, otherwise non-parametric comparisons were carried out. Environmental variables (O_2_ and T) were compared between seasons by Mann-Whitney *U* test. To analyze O_2_ and T heterogeneity we calculated the % coefficient of variation (% CV, SD/mean*100) for each census unit and then the mean %CV and SD per season. Fish individual traits (body length and EOD rate), and DNN were compared between sexes within each season by t-tests. Body length, EOD rate and DNN were compared seasonally by t-test for females and males together.

To examine the effect of individual traits (see below) on DNN as the dependent variable we used generalized linear models (GLM; (54)), in each season by separate. For the breeding season, we first ran a model with body length, EOD rate and sex as explanatory variables. Because the initial model was non-significant, we ran one model for females with body length, EOD rate, and circulating E_2_ as explanatory variables, and a second model for males with body length and EOD rate as explanatory variables. For the non-breeding season, the model included both, females and males together, and explanatory variables were body length, EOD rate and sex. For each season separately, initial models contained all single effects and pairwise interactions of the explanatory variables. To select the most parsimonious GLM, we used the command bestglm (55), for a maximum of 3 simultaneous variables, and considering up to second order interactions. Initial models were simplified by the stepwise deletion of the least significant terms in a model and comparing successive steps of model simplification by the Akaike information criterion (AIC), deleting a term whenever there was a difference of more than 2 units between alternative models until arriving to the most parsimonious model that could be fitted. The selection of the best model included the AIC criterion as well as the number and statistical significance of the estimated parameters, discarding models with improvements in the AIC but non-significant parameters. All models were subjected to the customary residual analysis (56).

To evaluate if the sex of the nearest neighbor was different from what is expected by a random distribution, we used Binomial tests (57). For the analysis, the focal fish was only considered if the nearest neighbor was retrieved and the sex determined. The proportion of sexes expected for a random distribution was deduced from the empirical sex ratio observed in each season.

Parametric, non-parametric statistical analyses, and simple linear regressions were carried out with PAST (58), and GLMs and Binomial tests with software R (59) using RStudio interface.

## Results

Individuals of *Gymnotus omarorum* were located at approximately 30 cm of depth among the dense roots of extensive floating mats of vegetation along the littoral area across seasons (Fig. 1B; i.e., in the breeding season during the austral summer from December to February, and in the non-breeding season during the austral winter from May to August). The water hyacinth *Eichhornia crassipes* dominated both surface and underwater areas accounting for 86% of the total subaquatic biomass. Associated vegetation was composed by the submerged *Egeria densa* and *Miriophyllum aquaticum*, the free-floating *Salvinia auriculata*, and the rooted but partly emergent *Ludwigia elegans*, and *Hydrocotyle criptocarpa*. This vegetation was present year-long although overall coverage was lower during the non-breeding season.

As expected in the subtropical region, water temperature and oxygen content showed significant differences across seasons. Water temperature was higher during the breeding than during the non-breeding season (27.3 ± 0.1 °C, N = 36 vs 11.3 ± 0.5 °C, N = 60; p < 0.0001, Mann Whitney *U* test), whereas O_2_ concentration was significantly lower in the breeding compared to the non-breeding season (5.6 ± 0.9 mg/l, N = 36, vs 8.4 ± 0.4 mg/l, N = 65; p = 0.004, Mann Whitney *U* test). In contrast, water conductivity remained consistently below 150 μS/cm throughout the year.

### Field survey 1: Fish spatial distribution and environmental variables

Fish (dots in Fig. 1C) were detected by electrical census in individual sites located under the central area of the floating mats, and absent from the edge limiting the open water (Fig. 1C). In both seasons, fish were found in an even distribution, non-aggregated with other conspecifics. The distribution of DNNs was asymmetrical, skewed with a mode at 1.5 m (Fig. 2A). The DNN mean value was significantly higher in the breeding season than in the non-breeding season (2.3 ± 0.15 m, N = 47 vs 1.4 ± 0.07 m, N = 73; p < 1 exp-4, t-test).

**Figure 2:**
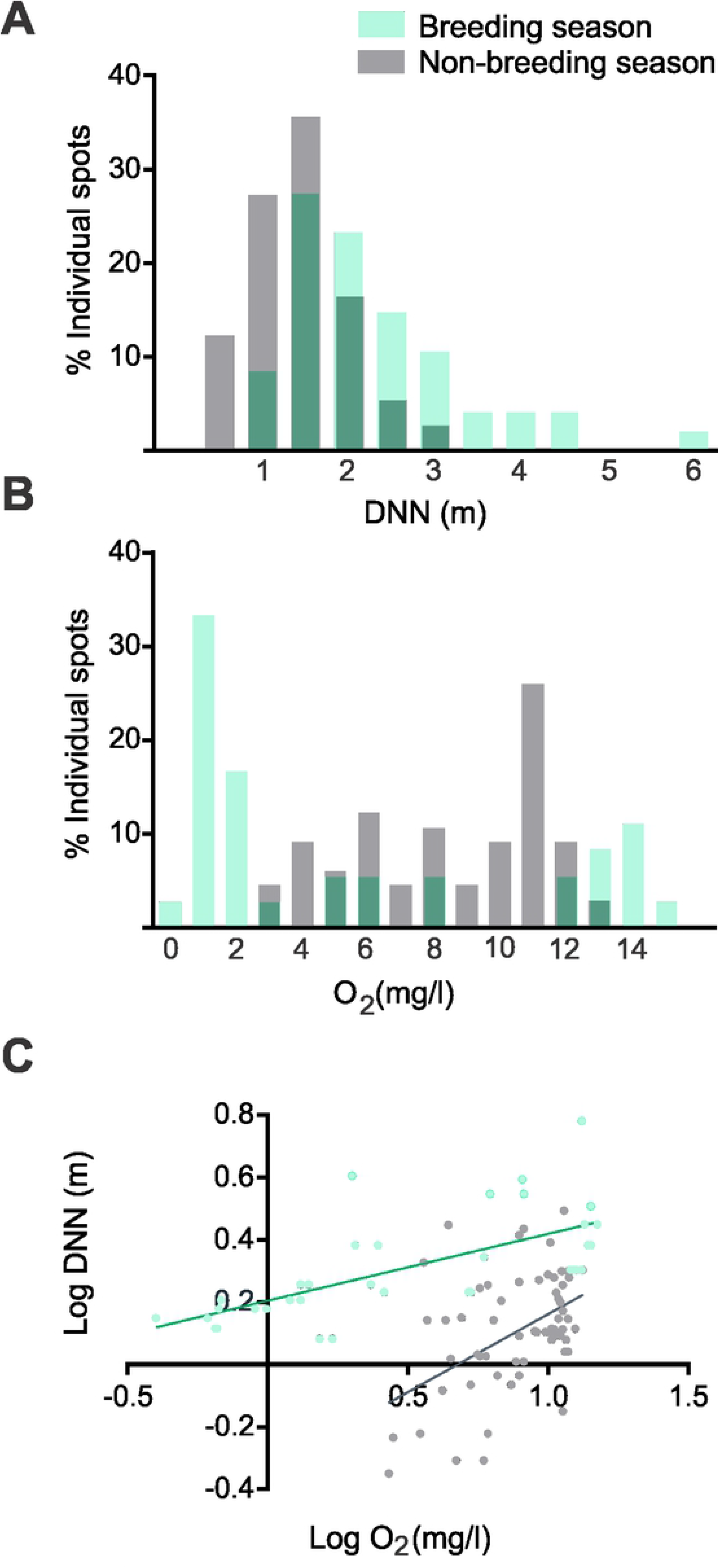
Fish spatial distribution based on environmental variables. The breeding season is represented in green and the non-breeding season in gray; dark green sections implies overlap of both seasons. A. Frequency distribution versus distances to the nearest neighbor (DNN, in meters). B. Frequency distribution versus oxygen concentration (in mg/l) measured at 30 cm from the surface in each individual spot. C. Linear correlation of distance to the nearest neighbor and oxygen concentration in individual spots Breeding season: R2 = 0.44, p = 1 exp −4, N = 31, non-breeding season: R2 = 0.21, p = 1 exp −4, N = 50.

Water temperature showed low variability among individual spots within each census unit (breeding season mean T CV = 2.0 ± 2.4 %, non-breeding season mean T CV = 6.5 ± 5.5 %). In contrast, O_2_ concentration showed high variability within each census unit (breeding season mean O_2_ CV = 20.4 ± 9.3 %, non-breeding season mean O_2_ CV = 25.9 ± 14.6 %), and thus we explored whether O_2_ could account for fish spatial patterns. During the breeding season, the distribution of O_2_ concentration ranged from 0 to 15 mg/l and was positively skewed (Fig. 2B), with a higher frequency of low values (mode at 1 mg/l). However, during the non-breeding season, O_2_ distribution was negatively skewed, showing a single mode at 11 mg/l (Fig. 2B). Oxygen concentration and DNN showed a positive and significant association both in the breeding season (R2 = 0.44, p = 1 exp −4, N = 31, Fig. 2C green) and the non-breeding season (R2 = 0.21, p = 1 exp −4, N = 50, Fig. 2C gray).

### Field survey 2: Fish spatial distribution based on individual traits

There were no sexual differences in body length, EOD rate or DNN in both the breeding and the non-breeding season (Table 1). During breeding, females showed circulating mean E_2_ levels of 293.7 ± 97.7 pg/ml and males had mean 11-KT circulating levels of 399.1 ± 140.9 pg/ml. From a seasonal perspective, breeding fish were significantly larger than non-breeding ones (p < 1 exp-4, N breeding season = 28, N non-breeding season = 53; t-test). In addition, EOD rate was higher (p < 1 exp-4, N breeding season = 30, N non-breeding season = 36; t-test) and DNN was larger during the breeding season (p < 1 exp-4, N breeding season = 31, N non-breeding season = 46, t-test).

**Table 1:** Sexual comparison of individual traits and distance to the nearest neighbor in the breeding season and in the non-breeding season. Individual traits: Body length and electric organ discharge (EOD) rate corrected by water temperature. Distance to the nearest neighbor (DNN). Values are expressed as mean ± standard error of the mean (SEM), and statistical comparisons carried out by t-test.

|  | Body size (cm) |  | EOD rate<sub>temp</sub> |  | DNN (m) |  |
| --- | --- | --- | --- | --- | --- | --- |
|  | ♀ | ♂ | ♀ | ♂ | ♀ | ♂ |
| **A- Breeding season** | 22.4 ± 0.5<br>n = 13 | 24.5 ± 0.9<br>n = 15 | 41.0 ± 1.1<br>n = 13 | 39.3 ± 0.9<br>n = 17 | 2.2 ± 0.2<br>n = 13 | 2.3 ± 0.3<br>n = 18 |
|  | ♀ vs ♂ p = 0.2 |  | ♀ vs ♂ p = 0.3 |  | ♀ vs ♂ p = 0.8 |  |
| B- Non-breeding season | 16.9 ± 0.9<br>n = 26 | 17.4 ± 0.8<br>n = 27 | 20.0 ± 1.2<br>n = 18 | 19.3 ± 0.95<br>n = 18 | 1.13 ± 0.1<br>n = 23 | 1.2 ± 0.1<br>n = 23 |
|  | ♀ vs ♂ p = 0.7 |  | ♀ vs ♂ p = 0.6 |  | ♀ vs ♂ p = 0.5 |  |

We evaluated the determinants of DNN separately in the breeding and non-breeding seasons. As the first GLM including breeding females and males, with body length, EOD rate, and sex as explanatory variables was not significant, we separated sexes into two different models and included circulating E_2_ levels as an explanatory variable for females.

We obtained two significant models in females, both equivalent according to the AIC criterion (Table 2). In the model with the best adjustment (model 1), DNN showed a positive correlation with circulating E_2_, and a negative correlation with EOD rate. In model 2, EOD rate was not a significant explanatory variable for DNN and only E_2_ had a significant positive correlation. For males, we were unable to find a correlation between the independent variables tested and DNN. Although androgen levels were not included in the model (due to a low number of valid samples), it is worth mentioning that circulating 11-KT levels showed a positive trend with DNN in a simple linear regression (p = 0.049; R2 = 0.77; N = 5). During the non-breeding season, as all traits quantified were represented in both sexes, we were able to run females and males together when testing the influence of individual traits on DNN. We explored if individual sex, body size, and EOD rate correlated with DNN, and found that body size, but not sex nor EOD rate, correlated positively with DNN (Table 3).

**Table 2:** GLM models that presented the best adjustment to explain the distance to the nearest neighbor in females during the breeding season. Model intercept and explanatory variables are expressed as Mean (SD). For each parameter the value is shown in bold if statistically significant (<0.05), in italic if marginal (<0.1), and expressed as NS if non-significant. Model 1 and Model 2 did not present significant differences by the Akaike information criterion (AIC).

| Model | Intercept | Circulating E<sub>2</sub> | EOD rate | N | P model | Adjusted R<sup>2</sup> | AIC |
| --- | --- | --- | --- | --- | --- | --- | --- |
| M1 | **4.52** (1.41) | **1.4e-3** (0.03) | *-0.051* (0.027) | 13 | 0.022 | 0.44 | 20.24 |
| M2 | **1.88** (0.18) | **1e-3** (4e-4) | NS | 13 | 0.028 | 0.31 | 22.21 |

**Table 3:** GLM model that presented the best adjustment to explain the distance to the nearest neighbor in both sexes during the non-breeding season. Model intercept and explanatory variable are expressed as Mean (SD). For each parameter the value is shown in bold if statistically significant (<0.05).

| Intercept | Body size | N | P model | Adjusted R<sup>2</sup> | AIC |
| --- | --- | --- | --- | --- | --- |
| 0.364 (0.34) | **0.052** (0.02) | 48 | 0.014 | 0.11 | 79.75 |

Although DNN was larger in the breeding season in both sexes, this seasonal difference disappeared in males when DNN was normalized by fish body size (0.08 ± 0.007, N = 17 vs 0.1 ± 0.01, N = 22; p = 0.16, t-test, Fig 3A). Interestingly, in females, DNN normalized by body size was significantly higher during the breeding season than in the non-breeding season (0.1 ± 0.008, N ; 13 vs 0.07 ± 0.006, N = 23; p = 0.01, t-test, Fig 3A).

**Figure 3:**
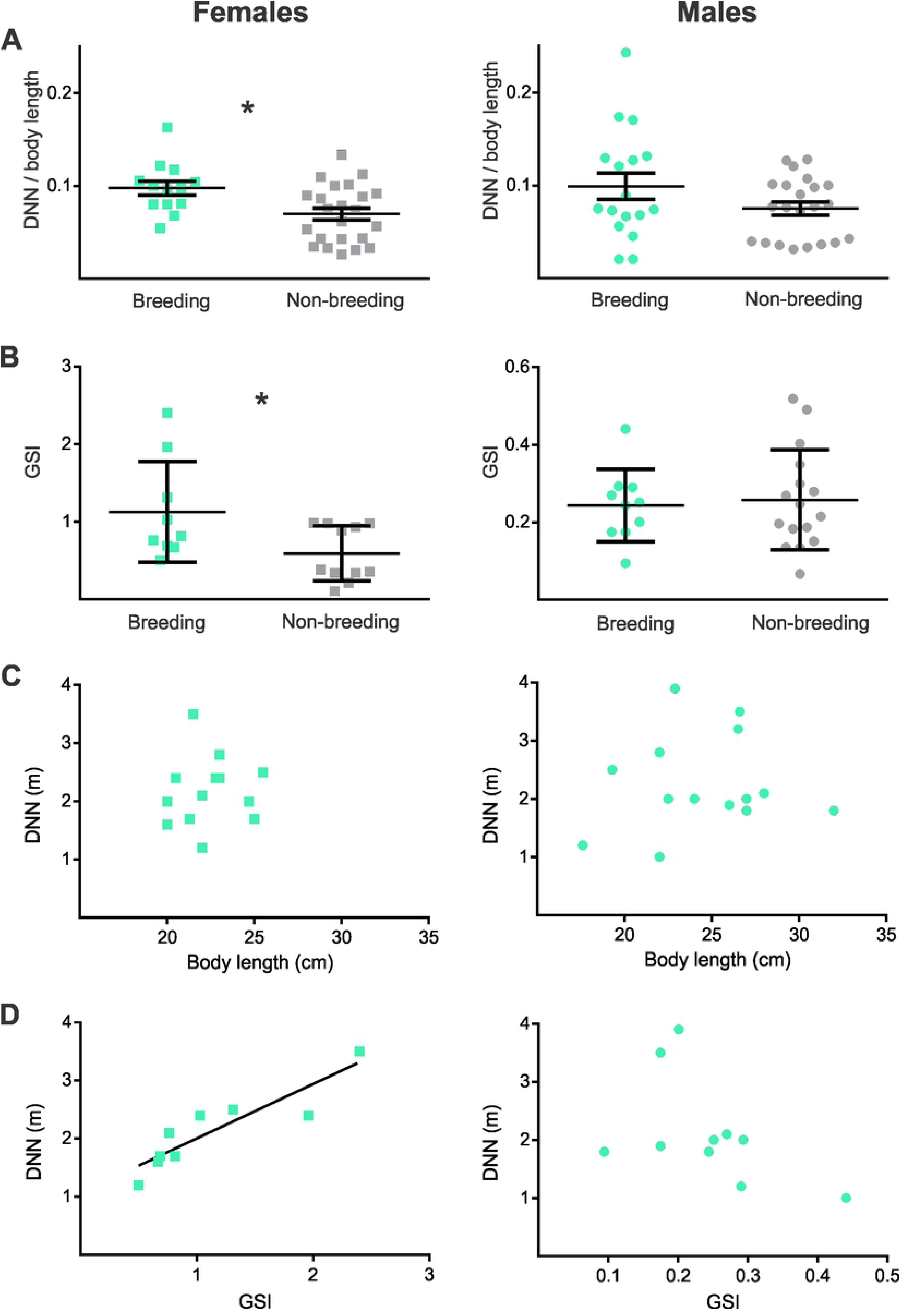
Emergence of sex dimorphism during the breeding season. The plots show values for females (left panels, represented as squares) and males (right panels, represented as circles); colors indicate breeding (green) and non-breeding (grey) seasons. **A-** Distance to the nearest neighbor (DNN in m) corrected by body length (cm). **B-** Gonadosomatic index (GSI). Dots represent individual values, and in A and B horizontal line represent mean values, and error bars represent SEM. For each sex, breeding values are shown in the left (green) and non-breeding values in the right (grey). * indicate statistically significance (p<0.05) t-test. **C-** Linear regression between body size (cm) and DNN (m) in the breeding season. Females: p = 0.8, R2 = 0.007, N = 13; males p = 0.7, R2 = 0.01, N = 15. **D-** Linear regression between GSI and DNN (m) in the breeding season. Females: p = 0.001, R2 = 0.8, N = 9; males: p = 0.14, R2 = 0.25, N = 10.

Animals sampled in both seasons differed in their gonadosomatic index depending on the sex (Fig. 3B). In females, GSI was significantly higher in the breeding season than in the non-breeding season (1.1 ± 0.22 % N = 9 vs 0.6 ± 0.09 % N = 11; p = 0.02, t-test), whereas males did not show seasonal differences (0.24 ± 0.03 % N = 10 vs 0.23 ± 0.03, N = 16; p = 0.87, t-test). Body size did not correlate with DNN in the breeding season (females, p = 0.8, R2 = 0.007, N = 13; males p = 0.7, R2 = 0.01, N = 15; Fig. 3C). In the breeding season, GSI strongly correlated with DNN in females (p = 0.001, R2 = 0.8, N = 9; Fig. 3D), whereas the correlation was not significant in males (p = 0.14, R2 = 0.25, N = 10, Fig. 3D).

In the breeding season, the spatial arrangement of sexes showed a specific configuration in which the percentage of females with a female as the nearest neighbor was significantly lower than the random distribution (p=0.03, N = 11; Binomial exact test, Fig. 4A). In the same sense, the percentage of males with a female as the nearest neighbor was significantly higher than the random distribution (p=0.04, N = 16; Binomial exact test, Fig. 4A). In contrast, the spatial configuration of the population in the non-breeding season showed a random arrangement of both males and females. The probability of having a female as the nearest neighbor was not significantly different from the random distribution for both females (p=0.82, N = 20; Binomial exact test, Fig. 4B) and males (p=0.12, N = 15; Binomial exact test, Fig. 4B).

**Figure 4:**
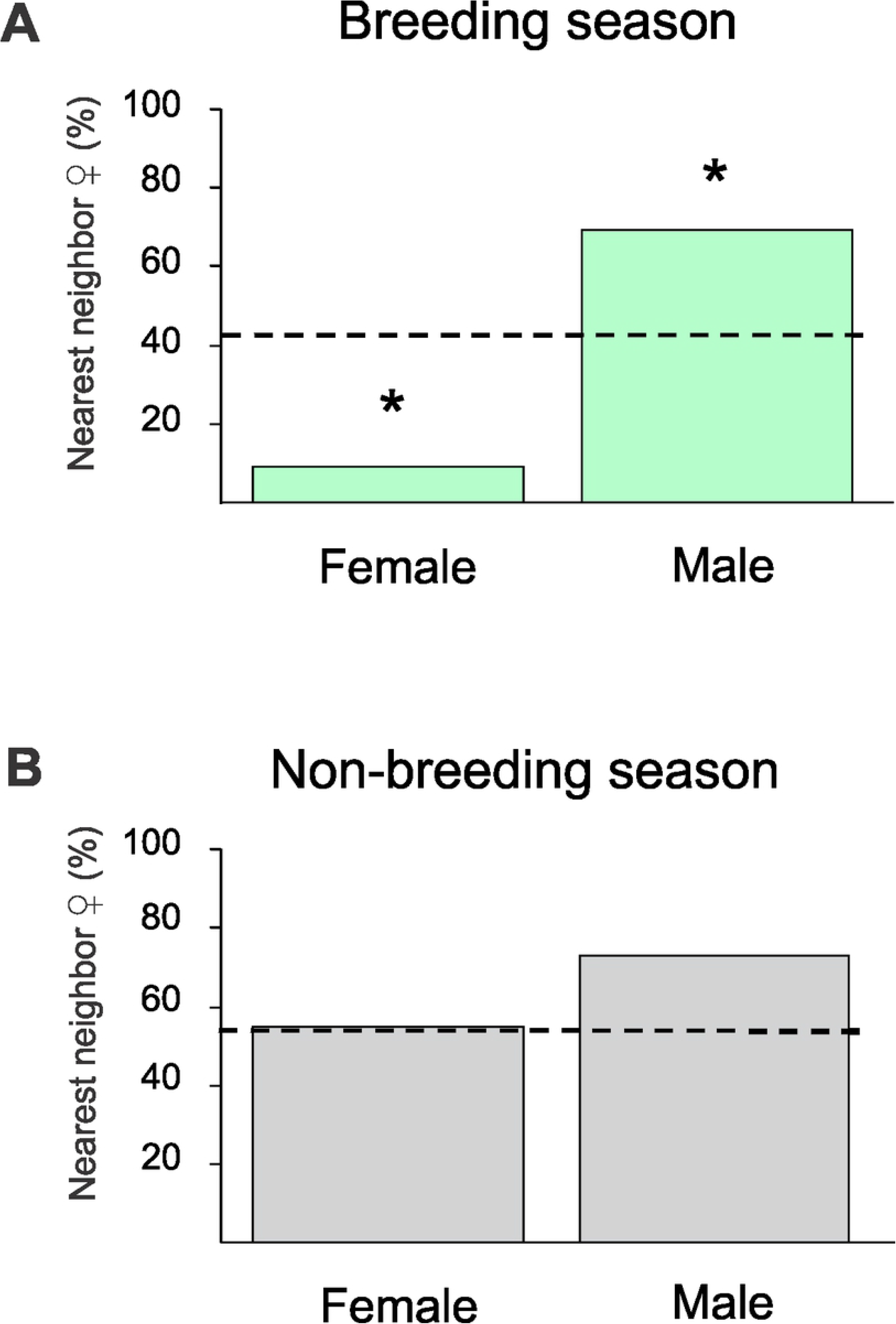
Seasonality in the spatial distribution of sexes. Sex of the nearest neighbor (expressed in percentage) when the focal fish is a female or a male, both in the breeding season (A, top), and in the non-breeding season (B, bottom). Dashed line represents the expected percentage to have a female as nearest neighbor for a random distribution, according to the empirical sex ratio (41% in the breeding season, and 50.9% in the non-breeding season). * indicate statistical significance (p<0.05) according to the binomial exact test.

## Discussion

This study contributes to the understanding of the mechanisms underlying behavioral plasticity by reporting for the first time seasonal and sexual differences in the determinants of territory size in a teleost species. In the first place, we found that *Gymnotus omaroum* presents a spatial arrangement in the natural habitat consistent with territoriality across seasons. In addition, we confirmed our predictions: a) In the non-breeding season, territory size was sexually monomorphic and partially determined by individual body size; and b) in the breeding season, sexual differences emerged given that the spatial arrangement adopted a sexual bias and that females hold gonadal-dependent larger territories.

Body mass has strong relationships with a variety of physiological and ecological attributes, such as home range and metabolic rates and is key to understanding how animals use the environment (60–63). The distribution of differently sized animals can be shaped by behavioral interactions such as the defense of foraging resources (64). These ideas, put forth initially to analyze interspecific interactions, can be also applied to interpret distribution of individuals of the same species, as in this study (65). We found that body size predicted DNN in the non-breeding season but not in the breeding season (Tables 2 and 3). This suggests that during reproduction other physiological, behavioral, and motivational aspects may be overriding body size as a predicting trait on DNN, a proxy for territory size.

Oxygen is a limiting physico-chemical variable in aquatic ecosystems, and low concentrations of dissolved O_2_ has been related to fish mortality (66). Interestingly, *Gymnotus* has been shown to be tolerant to a wide range of dissolved O_2_ concentration and can survive in places with hypoxia (67), as it is capable of breathing from air in addition to metabolic adaptations which compensate for any potential damages hypoxia may cause (68). Although different individual spots showed variable O_2_ concentration, and O_2_ correlated with territory size in both seasons, it remains to be understood whether high O_2_ concentration may be a valuable attribute of territories in this species. In this regard, despite being tolerant to hypoxia, higher concentration of available O_2_ could most probably offer a less demanding context for fish.

### Non-breeding spacing: body size dependent and sex independent

In teleosts, year-round territoriality has been mostly approached in populations of coral reef communities (for example *Stegastes fuscus*, (69)). Here, we evaluated the seasonal spatial organization in a subtropical freshwater fish in the field and found that spatial distribution during winter was consistent with territoriality. Why would *Gymnotus* defend non-breeding territories? The acquisition and maintenance of territories are known to be mediated by agonistic behavior (70). In particular, previous reports of laboratory experiments demonstrated that *Gymnotus omarorum* exhibits non-breeding agonistic behavior that mediates territory access (38) and that body size is the main determinant of the outcome of fights (22). Electric fish present high basal metabolic requirements associated to electrogeneration (71), which impose an additional constant need to forage. In line with this, we have found that larger fish hold larger territories (Table 3). Body size is sexually monomorphic in this species and behavioral experiments have shown that agonistic behavior is non-sex-biased (22,39). In this study, we confirmed that non-breeding males and females hold sexually monomorphic territory sizes in the wild, probably to cope with energetic requirements that are not expected to be sexually different during winter.

Electric fish use electric communication signals as behavioral displays. As EOD encodes information about body size and physiological state (72–74), territory boundaries could be maintained in the wild by remotely assessing the EOD of neighbors. Status dependent EOD rate-rank occurs only when fish are kept in close quarters after conflict resolution, but not when allowed to distance themselves (38). Thus, we did not expect to find correlations between EOD rate and territory size in the field, as was the case (Table 3). EOD rate rank may be a behavior needed to reinforce submission when a subordinate individual is unable to escape from the dominant as in confined laboratory conditions but not in the wild under the population densities observed in this study.

### Breeding spacing: the emergence of sexual dimorphism in territory determinants

While competition for reproductive opportunities is usually sexually dimorphic, competition over non-sexual resources can be expected to be balanced between males and females (75,76). Moreover, species of different phylogenetic groups with male and female territoriality have been found to display sexual monomorphism in body size and signal traits (77–79). Consistently, we found that during the breeding season absolute territory sizes were sexually monomorphic, which is consistent with the lack of sex dimorphism in body size (Table 1). However, during breeding, body size did not correlate with territory size (Fig. 3A and B), suggesting there are other factors overcoming body size influence on habitat resource acquisition and maintenance.

Interestingly, a closer analysis of our data showed that territory size relative to body size was sexually dimorphic. Females seemed to need larger territories in the breeding season compared to the non-breeding season (Fig. 3A), but males showed no seasonal difference (Fig. 3B). This result can be interpreted in the context of the energetic cost imposed by ovarian maturation, as GSI showed an excellent predicting power on female territory size (Fig. 3 A). Steroid hormones are crucial as mediators of behavioral plasticity (1). Sex steroids orchestrate integrated responses in the organism, and also respond depending on the social environment (80,81). Not surprisingly, circulating E_2_ positively correlated with territory size in females (Table 2). The fact that both GSI and E_2_ correlated with female territory size, in addition with reports in which E_2_ promotes female aggression (82–84), suggests that ovarian E_2_ modulates territorial behavior in *G. omarorum*. We hypothesize that E_2_ is integrating female metabolic requirements with the social environment, trough the expression of territorial behavior. On the other hand, males showed a trend between circulating 11-KT and territory size, which was expected given the well documented relationship between androgens and male territoriality (24–26).

In the breeding season, sexually dimorphic individual traits may influence motivation towards territory defense, and thus be reflected in the spatial pattern (11,12). The results presented in this study suggest that *G. omarorum* can assess territory features and use this information to relocate at different times of the year. In the non-breeding season individual fish had a closest neighbor which was randomly either of the same or opposite sex, whereas in the breeding season, it was more likely to have an opposite-sex closest neighbor (Fig. 4). This evidence supports the idea that the sex of the nearest neighbor becomes a relevant factor for territory value, but only during breeding. This can be considered a good example of behavioral plasticity by which individuals respond differently to the same social stimulus (e.g., sex of the nearest neighbor), depending on variations in their internal state (sexually dimorphic hormones).

### Concluding remarks

*Gymnotus omarorum* is a species which offers the opportunity to analyze seasonal changes in year-round territoriality. Although we found no differences in absolute territory size between sexes across the year, territory size was truly sexually monomorphic only in the non-breeding season, in which body size was the only significant determinant of territory size. In contrast, in the breeding season sex became relevant for the territorial behavior of this species: a) the sex of neighbors matters; b) territory size depended on gonadal hormones in both sexes, which was expected for males, but not previously reported in females; and c) females needed relatively larger territories than males which may reflect particular sex-biased reproductive requirements consistent with metabolic demands of anisogamy. This study contributes to bridge the gap between behavioral plasticity of natural territorial behavior and its underlying mechanisms.

## Author’s contributions

L.Z., L.Q., and A.S. conceived and designed the overall study.

M.M. and F.T.M. contributed to the design of field sampling.

L.Z., L.Q., F.T.M., M.M., and A.S. collected the data.

L.Z., R.H.M., and R.G.M. processed and interpreted hormone data.

L.Z. and D.H. analyzed all data.

L.Z., L.Q., D.H., F.T.M., M.M., and A.S. interpreted results.

L.Z., L.Q., and A.S. led the writing of the manuscript.

L.Q. and A.S. funding acquisition.

All authors contributed critically to the drafts and gave final approval for publication.

## Acknowledgements

We are very grateful to Adriana Migliaro, Rossana Perrone, Carlos Passos, Federico Reyes, and Bettina Tassino for their useful discussions during the BERTA Workshop, Cerro del Toro, Piriápolis, Uruguay. L.Z., L.Q., F.T.M., M.M., and A.S would like to thank to Agencia Nacional de Investigación e Innovación (ANII), Programa de Desarrollo de las Ciencias Básicas (PEDECIBA), and Universidad de la República, Uruguay for funding.

Specific grants that funded this research: ANII_FCE_6180, ANII_FCE_136381, FCE_4272.

